# 3D multi-color far-red single-molecule localization microscopy with probability-based fluorophore classification

**DOI:** 10.1101/2022.01.14.476290

**Authors:** Marijn E. Siemons, Daphne Jurriens, Carlas S. Smith, Lukas C. Kapitein

## Abstract

Single-Molecule Localization Microscopy remains limited in its ability for robust and simple multi-color imaging. Whereas the fluorophore Alexa647 is widely used due to its brightness and excellent blinking dynamics, other excellent blinking fluorophores, such as CF660 and CF680, spectrally overlap. Here we present Probability-based Fluorophore Classification, a method to perform multi-color SMLM with Alexa647, CF660 and CF680 that uses statistical decision theory for optimal classification. The emission is split in a short and long wavelength channel to enable classification and localization without a major loss in localization precision. Each emitter is classified using a Generalized Maximum Likelihood Ratio Test using the photon statistics of both channels. This easy-to-adopt approach does not require nanometer channel registration, is able to classify fluorophores with tunable low false positive rates (<0.5%) and optimal true positive rates and outperforms traditional ratiometric spectral de-mixing and Salvaged Fluorescence. We demonstrate its applicability on a variety of samples and targets.

## Introduction

15 years after its invention, Single-Molecule Localization Microscopy (SMLM) has developed into a reliable and widely used imaging modality to resolve structures beyond the diffraction limit [1-3]. The fluorophore Alexa647 (AF647) is the staple of the most popular SMLM technique called direct Stochastic Optical Reconstruction Microscopy (dSTORM) [4] due to its brightness and excellent blinking dynamics. However, finding spectrally complementary dyes for multi-color imaging has remained a challenge. It takes extensive tuning of laser power and buffers to optimize the blinking when using fluorophores outside of the far-red channel [5]. In contrast, other far-red fluorophores such as CF660 and CF680 also exhibit proper brightness and blinking, but display significant spectral overlap.

One way to overcome this challenge is the use of a grating or prism (spectroscopic SMLM) [6-8] or to encode the spectral information in the PSF [9]. However, these methods increase the footprint of the spot deteriorating the signal to background ratio and significantly increase the sparsity constraints, which makes them unsuitable for many applications. Another option is to use ratio-metric spectral de-mixing [10-12]. However, regular ratio-metric spectral de-mixing still requires significant separated emission spectra (i.e. AF647 and CF660 cannot be used without major crosstalk or significant rejection). Another complication of ratiometric spectral de-mixing is that it requires nanometer registration of the imaging channels. In order to perform this correctly, chromatic aberrations and field distortions have to be calibrated with a high precision, about 20 to 50 times smaller than the pixel size, to ensure super-resolution reconstructions without significant misalignment [10]. Therefore artefacts can be easily introduced when calibration is not performed correctly and frequently.

Recently, an alternative way to spectrally de-mix AF647 and CF660 was demonstrated on a 4Pi microscope, in an approach termed ‘salvaged fluorescence’ detection [13]. Here localization and detection is performed using the fluorescence collected in the regular imaging channel, but the fluorescence reflected by the dichroic mirror that couples in the excitation light (called ‘salvaged fluorescence’) is used for classification. This captures the low wavelength front of the emission spectrum, which is the most distinguishable feature of the different far-red fluorophores. As such, this small wavelength window enables adequate classification without compromising the detection and localization in the other channel. Furthermore, this method does not require nanometer channel registration. However, the proposed approach to estimate the salvaged fluorescence includes the background level and is therefore sensitive to experimental changes that affect this, such as the chosen labeling targets or exposure time. In addition, in conventional microscopes it is challenging to detect the light reflected by the excitation dichroic mirror, which has so far limited the implementation of the approach in other systems.

Here we present the multi-color SMLM approach PFC (Probability-based Fluorophore Classification), which is implementable on conventional microscopes, does not require nanometer channel registration and enables three-color imaging of AF647, CF660 and CF680 with minimal crosstalk. Inspired by the salvaged fluorescence concept, the emission is split in a high intensity, long wavelength channel, used for detection and localization, and a low intensity, short wavelength channel used to facilitate classification. However, by optimizing the choice of dichroic mirrors and filters both channels are now imaged on a single camera. Furthermore, classification is performed using both channels with a statistical test called a Generalized Likelihood Ratio Test (GLRT) [14]. Such a test has been demonstrated to distinguish optimally between random background fluctuations and (dim) single-molecule blinking events [15]. In our case, the GLRT can determine the most likely fluorophore candidate for the blinking event, given the measured pixel values in both channels. This novel spectral de-mixing method allows for the classification between the spectrally very close fluorophores AF647, CF660 and CF680, which cannot be classified with traditional ratiometric spectral de-mixing or Salvaged Fluorescence without rejecting a large number of detected fluorophores. We demonstrate this method for 2-color dSTORM (with AF647 and CF660 or CF680) and 3-color dSTORM (with AF647, CF660 and CF680) in both 2D and 3D using astigmatic PSF engineering [16].

## Results

### Setup

We used a regular TIRF microscope equipped with a dual channel module and chose our filters in such a way that all the fluorescence is collected and split onto a single camera (see Material and Methods). The emission was split in a short channel (channel 1) with intensity fraction 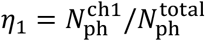 and a long channel (channel 2) with fraction *η*_*2*_ = (1 − *η*1). See Supplementary Figure 1 for the spectral characteristics of all the components. These spectral dichroic mirrors and filters were chosen such that the first part of the emission peak of AF647 was just captured in channel 1, resulting in intensity fractions of 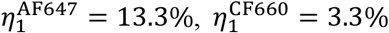 and 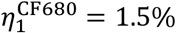 for AF647, CF660 and CF680, respectively (see Figure 1 and Supplementary Figure 2). Detection and localization was performed in the long channel which collects 86.7%, 96.7% and 98.5% of their fluorescence, respectively. For a 500 photon event, the small loss in intensity induced by this separation corresponds to a drop in localization precision of roughly 1 nm, 0.2 nm and 0.1 nm in the case for AF647, CF660 and CF680 respectively.

**Figure 1.**
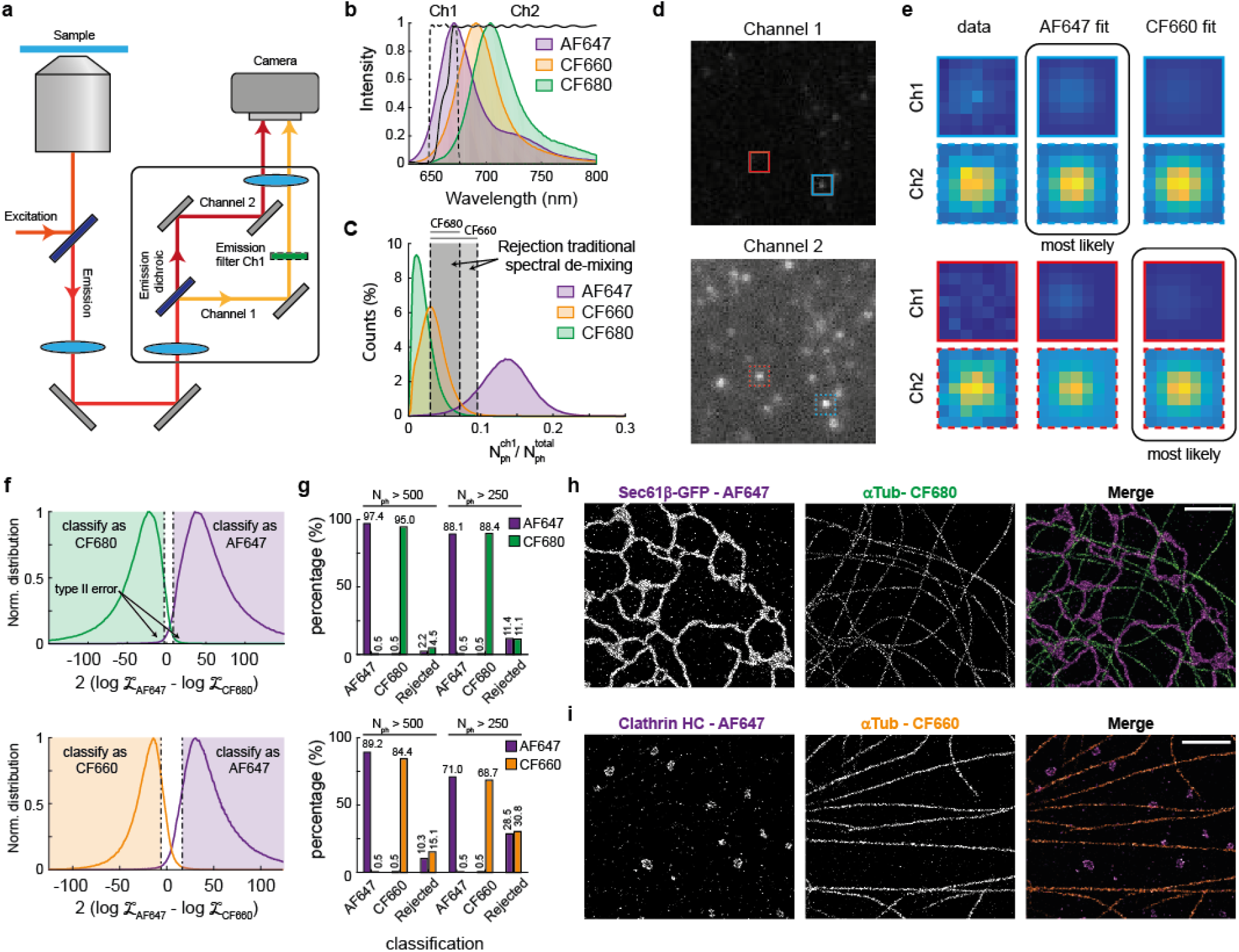
Two-color SMLM using PFC-dSTORM. a) Simplified diagram of the setup. b) Emission spectra of AF647, CF660 and CF680, overlayed with the emission dichroic (solid line) and channel 1 emission filter (dashed line). c) Distribution of the measured intensity fraction of channel 1 for events with 500 photons or more (n = 9.5×10^5^ events for AF647, n = 7.8×10^5^ events for CF660 and n = 9.6×10^5^ for CF680, N=5 acquisitions for each fluorophore). Grey regions indicate rejection areas when using traditional spectral de-mixing to achieve a false positive rate of 0.5%. d) Example acquisition of channel 1 and channel 2 with a sample labeled with AF647 and CF660. e) Example of the GLRT classification and the 2 MLE fits. f) Distribution of the GLRT with AF647 versus CF680 (top) and AF647 versus CF660 (bottom) for all events of c. Dashed lines indicate the cutoff values. Events with a GLRT value between the cutoffs are rejected. g) Classification percentages for all events with photon counts 500 and 250 or more for AF647 versus CF680 (top) and AF647 versus CF660 (bottom). h) Example 2-color PFC-dSTORM reconstruction of a COS-7 cell stained for ER (Sec61b-GFP overexpression, magenta) and alpha-tubulin (green) using AF647 and CF680, respectively. Scale bar indicates 2 μm. i) Example 2-color PFC-dSTORM reconstruction of a COS-7 cell stained for clathrin HC (magenta) and alpha-tubulin (orange) with AF647 and CF660. Scale bar indicates 2 μm.

### Generalized Likelihood Ratio Test for fluorophore classification

The Generalized Likelihood Ratio Test can classify the fluorophores based on the prior knowledge that a specific blinking event is either caused by fluorophore A or fluorophore B, which will yield two different intensity ratios between channel 1 and 2. The GLRT therefore has to test the following hypotheses

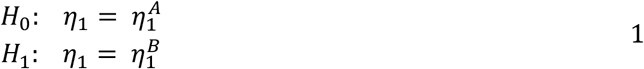

with 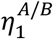 the (calibrated) intensity fraction in channel 1 for fluorophore A or B. This leads to the test statistic *T*, given by

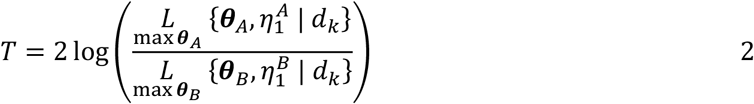

Where 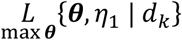 denotes the maximum likelihood obtained by a 2-channel MLE fit of pixel data *d*_*k*_ with fit parameters ***θ*** and fixed intensity fraction *η*1. This MLE fit procedure fits two coupled Gaussian distributions to the two spots, where the *η*1 governs the intensity ratio between the two Gaussian distributions (see Supplementary Note for details). To obtain the test statistic value, the two spots of a single blinking event are fitted twice: once with a fixed intensity fraction 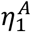 (assuming it is fluorophore A) and once with a fixed intensity fraction 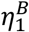 (assuming it is fluorophore B, see Figure 1d). The GLRT, which determines which fluorophore is the most likely candidate for a blinking event, provides the decision rule

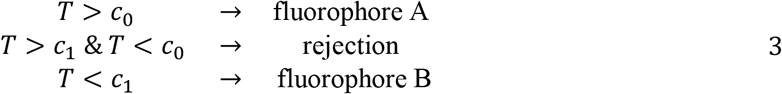

with *c*_0/1_ adjustable thresholds. These thresholds can be chosen to reduce the false positive rates, P(*T* > *c*_0_|*H*_1_) and P(*T* < *c*_1_|*H*_0_), and achieve a significance level α via P(*T* > *c*_0_|*H*_0_) = α and P(*T* < *c*_1_|*H*_1_) = α. As stated by the Neyman–Pearson lemma [14], this likelihood-ratio test is the most powerful among all level-α tests and can therefore classify the fluorophores with the lowest possible false positive rate for a chosen threshold *c*_*i*_. In the case for 3 or more fluorophores, a possible implementation is to test which of the models is the most likely [17]. However, here we perform the GLRT recursively (fluorophore A vs B followed by fluorophore B vs C) which is possible because 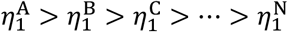. This allows for multiple thresholds to tune the false positive rates of each fluorophore.

### Classification performance

We first analyzed the performance of the PFC for dual color cases (AF647 vs CF660 and AF647 vs CF680). We experimentally obtained the distributions of P(*T*|*H*_*i*_) by measuring the values of the test statistic of blinking events in samples labeled with a single fluorophore (see Figure 1f). We preferred this experimental approach because it captures the natural variance in the intensity fraction (no event will have the exact calibrated intensity fraction) and it also includes possible SMLM imperfections, such as overlapping events or other blinking artefacts. This approach therefore gives a realistic false positive rate. We chose cutoff values of *c*_0_ = 9 and *c*_1_ = −3 to achieve false positive rates of 0.5% for both AF647 and CF680. This resulted in successfully classified fractions of 97.4% and 95% of the events as AF647 and CF680, respectively, with unclassified fractions of only 2.2% and 4.5% when considering events with 500 photons or more (see Figure 1g and Supplementary Figure 3). In traditional ratiometric de-mixing, 5.4% and 21.3% would have to be rejected for AF647 and CF680 in order to achieve identical false positive rates. The distributions of P(*T*|*H*_*i*_) can be approximated by two Gaussians when binned for photon count and the distance between these two Gaussians increased for higher photon counts (see Supplementary Figure 4). For this reason, less stringent cutoffs could be used for events with higher photon counts, which would result in a lower rejection rate of these high-intensity events, but a larger total amount of rejected events. A similar classification performance was achieved for AF647 in combination with CF660. In this combination, 10.3% and 15.1% has to be rejected respectively in order to achieve false positivity rates of 0.5%. This is again significantly lower than the rejected fraction in the case of traditional ratiometric spectral de-mixing in this configuration (16.1% and 55.6% respectively).

We next compared the performance of PFC to the classification scheme used in Salvaged Fluorescence. The Salvaged Fluorescence metric integrates the camera signal of the spot in channel 1 multiplied by a Gaussian mask and thereby does not distinguish between signal and background. The metric is therefore biased; it favors CF660 and CF680 in low background conditions and AF647 in high background conditions. This could be problematic because the background intensities might differ when using different labeling targets, sample preparation or imaging conditions such as exposure time. In contrast, the PFC algorithm includes the background in channel 1 as a separate fit parameter, which prevents biases when background intensities differ from the calibration condition. For our comparison, we again used the single-fluorophore samples to determine, for the desired false-positive rate of 0.5%, the rejection rates for different fluorophore combinations. It should be noted that this overestimates the performance of SF, because in these single-fluorophore samples the background in channel 1 is lower for CF660 and CF680 than for AF647. In these SF-biased conditions, SF achieves a similar classification performance as PFC for CF660 or CF680 (SF: 25.1% and 11.1% rejection for a photon threshold of 250, PFC: 29.1% and 9.5% rejection, for conditions where AF647 is the second fluorophore, Supplementary Figure 3). However, for AF647 PFC strongly outperformed SF. While SF rejects 74.5% and 42.9% of AF647 events in conditions with CF660 or CF680 as the second fluorophore, respectively, PFC only rejects 28.8% and 12.2%, respectively. These results demonstrates that our probability-based classification approach outperforms both traditional ratiometric de-mixing as well as Salvaged Fluorescence.

With our method we were able to perform 2-color dSTORM with both fluorophore combinations (see Figure 1h&i). We observed a clean separation between ER, labelled with AF647, and microtubules labeled with CF680 (see Figure 1h). We furthermore observed clearly visible clathrin coated vesicles and pits alongside densely labeled microtubules with no noticeable crosstalk using AF647 and CF660 (see Figure 1i). To illustrate the wide applicability of this method we show a collection of our multi-color imaging modality for a variety of targets in Supplementary Figure 5&6 (i.e. different microtubule subsets, microtubules and mitochondria, pre-and postsynaptic markers).

### 3-color imaging

We next tested if we could extend our approach to 3-color imaging. To achieve sufficient separation in the test statistics we introduced a different emission dichroic mirror (see Figure 2a and Supplementary Figure 1). The intensity fractions in channel 1 are in this case 27%, 8% and 3% for AF647, CF660 and CF680, respectively (see Figure 2b). The overlap between the intensity fraction distribution, shown in Figure 2b, clearly shows that traditional ratiometric spectral de-mixing cannot be used to separate CF660 and CF680, because it would entail that all events of CF680 need to be rejected. Although correctly classifying these events seems a daunting task, PFC is able to classify these event with acceptable false positive and rejection rates. To accomplish this, each blinking event is tested for AF647 vs CF660 and CF660 vs CF680. The distribution of the test statistics *T*_*A*F647 vs CF660_ and *T*_CF660 vs CF680_ can then be plotted in a 2D histogram, where each quadrant is associated with a unique fluorophore or rejection (see Figure 2c). Again, appropriate cutoff values for classification can be introduced to achieve the desired false positive rates (Figure 2d&e). In this case, false positive rates of 1% can be achieved while rejecting 0.1% of AF647, 28.5% of CF660 and 38.6% of CF680 for events which emitted 500 photons or more.

**Figure 2.**
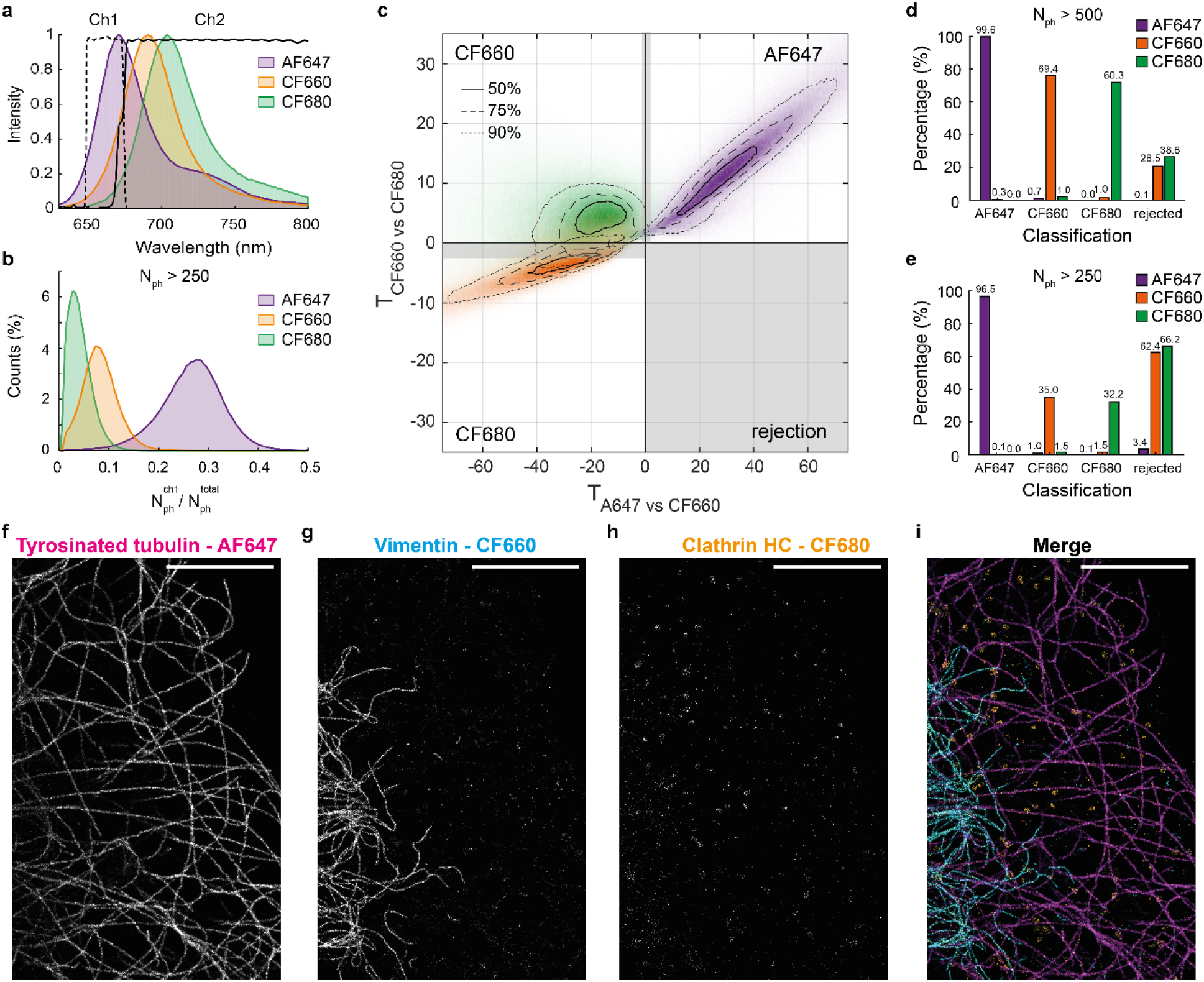
Three-Color SMLM with PFC-dSTORM. a) Emission spectra of AF647, CF660 and CF680, overlaid with the emission dichroic (solid line) and channel 1 emission filter (dashed line). b) Distribution of the measured intensity fraction of channel 1 for events with 250 photons or more (n=7.3e5 events for AF647, n = 1.1e6 events for CF660 and n = 5.1e5 for CF680, N=5 acquisitions for each fluorophore). c) 2D histogram of the test statistics AF647 versus CF660 and CF660 versus CF680 for the events shown in b. Grey area indicates rejection zone. (Dotted) lines indicate regions containing 50%, 75% and 90%, as indicated. d&e) Classification rates for photon thresholds of 500 and 250. f) Example 3 color PFC-dSTORM reconstruction of a COS-7 cell stained for tyrosinated tubulin (magenta), vimentin (cyan) and clathrin heavy chain (orange) with AF647, CF660 and CF680 respectively. Scale bar indicates 5 μm.

To demonstrate the 3-color capabilities of PFC in dense and overlapping structures we stained COS-7 cells for tyrosinated tubulin, vimentin and clathrin heavy chain (Figure 2f-i). We observed a clear separation between the microtubule network, the intermediate filaments and the clathrin coated pits. However, there appeared to be some crosstalk from the CF660 channel to the CF680 channel at sites where vimentin is abundant. This is expected when there are large discrepancies in the abundance of the stained structure, even with low false positive rates. Reconstructions of the full field-of-view are shown in Supplementary Figure 7.

### 3D imaging

Finally, we extended our multi-color SMLM approach to 3D localization by using astigmatic PSF engineering using a cylindrical lens module. For this, we modified the 2-channel MLE fit required for the GLRT to fit asymmetric Gaussians, which introduced an additional fit parameter (see Supplementary Notes for details). We performed 2-color 3D SMLM with astigmatic PSF engineering on COS-7 cells stained for ER and microtubules (see Figure 3). Our method was able resolve a microtubule width of 40 nm, consistent with immunolabeling [18] (see Figure 3d&g). Furthermore we were able to resolve the nanoscale ER morphology and observed ER matrices, consistent with recent findings using super-resolution microscopy [19] (see Figure 3e). Additionally, we found ER tubulation in the cellular periphery directly adjacent to microtubules, likely as result of microtubule dependent ER remodeling [20] (see Figure 3f). Lastly, our imaging modality was able to resolve hollow ER tubules in 3D at certain locations (see Figure 3i). Altogether, this shows that PFC allows the study of ER -cytoskeleton interaction with nanometer resolution in 3D.

**Figure 3.**
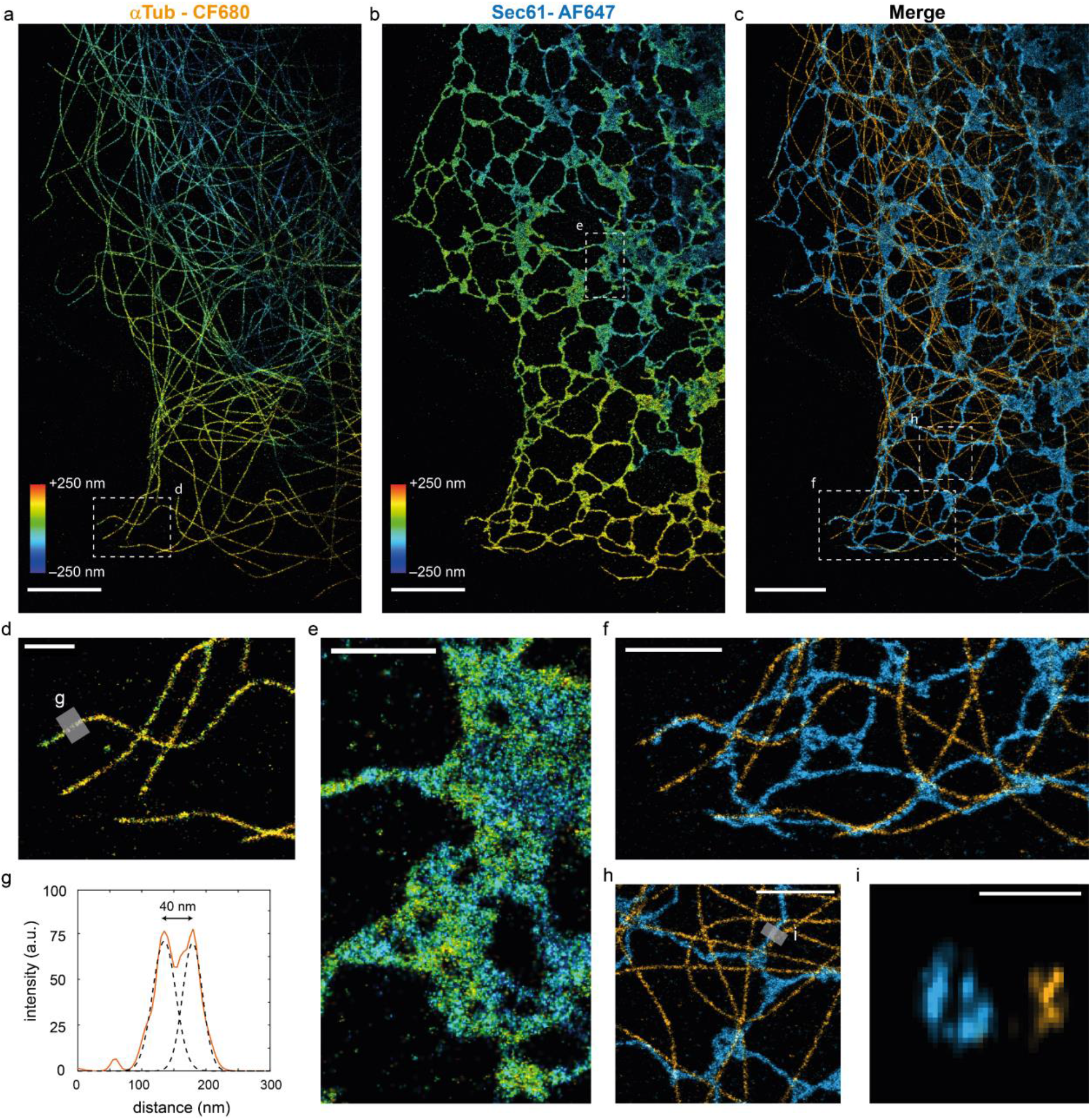
3D multicolor SMLM with PFC-dSTORM. a-c) Example 2-color 3D PFC-dSTORM reconstruction of a COS-7 cell stained for ER (Sec61-GFP overexpression, cyan) and alpha-tubulin (orange) with AF647 and CF680, color-coded for depth. d-f) Zooms of a, b and c. g) Intensity distribution of region indicated at d. h) Zoom of c. i) Cross section of region indicated at h. Scalebars indicates 5 μm (a, b, c), 2 μm (d, e, f, h) and 200 nm (i).

## Discussion

In this work we introduced a new method for multi-color SMLM, termed probability-based fluorophore classification (PFC), which featured two innovations over earlier work. Firstly, the emission is split in a high intensity channel and a low intensity channel, inspired by the salvaged fluorescence approach. However, our implementation only requires a single camera, does not require major rework on the microscope and is universally implementable. This approach allowed us perform the nanometric localization on just a single channel, which minimizes chromatic aberrations. The small associated loss in localization precision due to the photon loss in channel 1 mostly affects AF647, which is mitigated by the fact that AF647 is one of the brightest fluorophores available. Secondly, we introduced a Generalized Likelihood Ratio Test for fluorophore classification and implemented this test to be insensitive for channel misalignments. Therefore only a course pixel-to-pixel channel registration is required. This makes PFC a robust and convenient method that can be implemented by any lab with a TIRF microscope. Furthermore, the GLRT outperforms traditional ratiometric spectral de-mixing and Salvaged Fluorescence and can classify fluorophores with false positive rates as low as 0.5% with optimal rejection. This allows the use of the 3 best far-red fluorophores for dSTORM for imaging up to 3 colors. In the 3-color configuration AF647 can be classified without almost any rejection, while there is some rejection required for CF680 and CF660 in order to reduce crosstalk. This can potentially be overcome by choosing a slightly red-shifted emission dichroic, at the cost of a decrease in the intensity available for localization of AF647. Nonetheless, compared to spectral imaging modalities that use prims or gratings [6-8], this method is very photon-efficient, as these other methods often require 50% or more of the fluorescence for wavelength estimation.

This method also has several advantages over other multi-color SMLM approaches such as multiplexed DNA-PAINT [21], where a single acquisition can take multiple hours and which requires components with a very short shelf lifetime. Lastly, our implementation has no (additional) sparsity constraints compared to other methods using PSF engineering [9] or other ‘2-spot’ modalities [22, 23]. Therefore, dense structures such as microtubules and ER can still be imaged simultaneously. We demonstrated PFC dSTORM on a variety of samples and structures, such as the cytoskeleton network, ER and neuronal synapses, both in 2D and 3D, and show that this method is compatible with many different cellular components and is able to separate these with minimal crosstalk. We therefore anticipate this approach to become the go-to method for multi-color SMLM.

## Methods

### Setup

The setup consisted of a Nikon TI-E microscope equipped with a TIRF APO objective lens (NA = 1.49, 100X). A 638 nm laser (MM, 500mW, Omicron) was used for TIRF excitation via a laser clean-up filter (LL01-638, Semrock) and excitation dichroic (FF649-Di01, Semrock). The collected emission was filtered by a emission filter (BLP01-633R-25, Semrock) and relayed via a 1.5X tube lens (2D imaging) or 1X tube lens (3D imaging) to the emission port equipped with a cylindrical lens module (Nikon) and an Optosplit III module (Cairn Research). The emission dichroic (FF660-Di02, Semrock for 2 color imaging and Di03-R660-t1, Semrock for 3 color imaging) splitted the emission in a short channel and a long channel on a EMCCD (iXon 897 – Andor). See the Supplementary Note for details on the calibration. An additional emission filter (FF01-661/20-25, Semrock) was placed in channel 1. See Supplementary Figure 1 for the corresponding spectral characteristics of all the components.

### PSF-model and 2-channel MLE fit

We modeled the PSF in each channel as a, a simplification of the model used in [24], where the intensity *μ*_*k*_,_*λ*_ at pixel location _*k*_ for the respective channel *λ* is given by

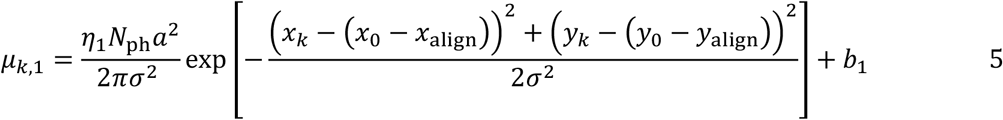

And

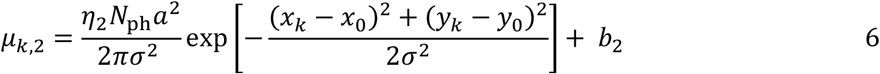

with *N*_ph_ the total number of emitted photons, *a* the pixelsize, *x*_0_ and *y*_0_ the position of the molecule, *x*_/_*y*_align_ a possible subpixel alignment correction between the channels, *σ* the width of both Gaussian PSFs and *b*_1/2_ the background in each channel. The log likelihood of the fit [9] is given by

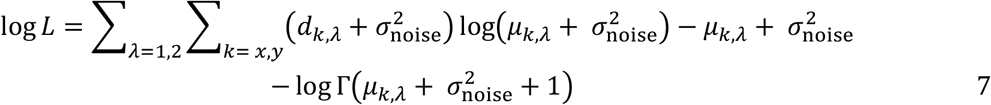

with *d*_*k*_,_*λ*_ the observed value of pixel *k* in channel *λ* and *σ*_noise_ the read noise of the camera pixel, which we assume to be zero for the EMCCD. The two spots in channel 1 and 2 are fitted simultaneously, leading to 8 fit parameters (***θ*** = *x*_0_, *y*_0_, *x*_align_, *y*_align_, *σ, N*_ph_, *b*_1_, *b*_*2*_) in total. In the case for astigmatic PSF a *x*- and *y*-directional width is fitted. See Supplementary Note on details of the fit algorithm.

### Intensity calibration

The intensity fractions for each fluorophore is calibrated by imaging COS-7 cells stained for αTub with a single fluorophore. The intensity in channel 2 is estimated with a regular 2D Gauss MLE fit which fits the x/y-position, width, intensity and background. The intensity in channel 1 is difficult to estimate as these photon counts are extremely low compared to the background level. To overcome this calibration issue we fit each spot in channel 1 with a 2D Gauss with a fixed width, obtained from the estimated width of the high intensity spot in channel 2. Lastly, fits are classified as outlier and removed if the log likelihood is smaller than the average log likelihood minus 3 standard deviations or if the estimated photon count is below 3 (channel 1) or 100 (channel 2). The intensity fraction is then estimated from the estimated photon counts in each channel of all spots with a weighted least-squares linear fit, where the weight is taken as the square root of the total estimated photon count of each spot.

### Sample preparation

#### Animals

In this study female pregnant Wistar rats were obtained from Janvier, and embryos (both genders) at E18 stage of development were used for primary cultures of hippocampal neurons. All experiments were approved by the DEC Dutch Animal Experiments Committee (Dier Experimenten Commissie), performed in line with institutional guidelines of University Utrecht, and conducted in agreement with Dutch law (Wet op de Dierproeven, 1996) and European regulations (Directive 2010/63/EU).

#### Cell culture

COS-7 and U2OS cells were grown in DMEM (Lonza, 12-604F) supplemented with 10% fetal calf serum (FCS, Sigma, F7524) at 37°C with 5% CO_2_. Dissociated hippocampal neuron cultures were prepared from rat pups at embryonic day 18 as described previously [25]. Briefly, cells were plated on 18-mm glass coverslips coated with laminin (1.25 mg/ml) and poly-L-lysine (37.5 mg/ml)(P8920 Sigma Aldrich) at a 50K/well density. Cells were maintained in Neurobasal medium (NB, Gibco, 21103-049) supplemented with 2% B27 (Gibco, 17504001), 0.5 mM glutamine (Gibco, 25030-032), 15.6 μM glutamic acid, and 1% penicillin/streptomycin (Sigma, P0781) at 37°C in 5% CO_2_.

#### Plasmids and transfection

For visualizing the ER we overexpressed GFP-Sec61β (Addgene #15108), an ER membrane protein. For transfection, DNA (1 μg) was mixed with 3 μl Fugene6 (Roche, #11836145001) in 200 μl opti-MEM (Gibco, 31985-047) and added to the cells for 16 hours or until fixation at 37°C with 5% CO_2_.

#### Fixation

Depending on the different structures that were targeted, three different fixation protocols were used: pre-extraction protocol, glutaraldehyde fixation protocol and PFA fixation protocol. For samples to be labeled for Tubulin, Clathrin HC and Vimentin we used the pre-extraction protocol, for samples with Sec61b-GFP overexpression we used the glutaraldehyde fixation protocol and for the samples to be labeled for Cytochrome C we used the PFA fixation protocol. All are described below.

The pre-extraction protocol was used for most cytoskeletal structures to remove the cytosolic pool of monomers. Cells were pre-extracted for 1 minute in extraction buffer (0.3% Triton X-100 (Sigma X100), 0.1% glutaraldehyde (GA) (Sigma G7526) in MRB80 buffer (80 mM Pipes (Sigma P1851), 1 mM EGTA (Sigma E4378), 4 mM MgCl_2_, pH 6.8), pre-warmed at 37°C. Afterwards, cells were fixed for 10 minutes in 4% EM-grade paraformaldehyde (PFA) (Electron Microscopy Science, 15710) and 4% sucrose in MRB80 buffer (pre-warmed at 37°C).

When targeting membrane bound structures the pre-extraction protocol cannot be used as this dissolves the membranes before fixation. We therefore used an alternative protocol that uses GA and PFA in cytoskeleton preserving buffer. Cells are fixed using 0.1% GA, 4% PFA and 4% sucrose in MRB80 buffer for 10 minutes (pre-warmed at 37°C).

Unfortunately, not all antibodies are compatible with glutaraldehyde, which results in a loss of signal intensity. For Cytochrome C we therefore fixed cells using 4% PFA and 4% sucrose in MRB80 buffer for 10 minutes (pre-warmed at 37°C).

#### Immunostaining

After fixation cells were washed 3 times in PBS (1 quick wash, followed by 2 washes of 5 minute) and permeabilized for 10 minutes with 0.25% Triton-X in MRB80. After again washing 3 times with PBS samples were further incubated for 1 hour in blocking buffer (3% w/v BSA in MRB80 buffer) at room temperature. Next, samples were incubated overnight at 4°C in primary antibodies diluted in blocking buffer. To proceed cells were washed 3 times in PBS before incubating for 1 hour at room temperature with secondary antibodies diluted in blocking buffer. After incubation cells were once more washed 3 times in PBS and kept in PBS at 4°C or mounted for imaging.

**Table 1.** Primary antibodies

| Target protein | Species | Dilution | Supplier | Cat # | Clone | Lot # |
| --- | --- | --- | --- | --- | --- | --- |
| clathrin heavy chain | mouse | 1/500 | Thermo fisher | MA1-065 | X22 | VL315162 |
| α-tubulin | mouse | 1/1000 | Sigma | T5168 | B-5-1-2 | 047M4760V |
| α-tubulin | rabbit | 1/1000 | Abcam | 52866 | EP1332Y | GR3241328-2 |
| gfp | chicken | 1/1000 | Aves Lab | GFP1010 | polyclonal | GFP3717982 |
| vimentin | rabbit | 1/300 | Abcam | ab92547 | EPR3776 | GR3258719-5 |
| tyrosinated tubulin | rat | 1/250 | Abcam | ab6160 | YL1/2 | GR3377281-5 |
| acetylated tubulin | mouse | 1/600 | Sigma | T7451 | 6-11B-1 | 059M4812V |
| homer | rabbit | 1/600 | SySy | 160 002 | polyclonal | Gift from<br>Hoogenraad lab |
| bassoon | mouse | 1/600 | Enzo | ADI-VAM-PS003-F | SAP7F407 | 06231712 |

**Table 2.** Secondary antibodies

| Host species | Target species | Fluorophore | Dilution | Supplier | Cat # | Lot # |
| --- | --- | --- | --- | --- | --- | --- |
| goat | chicken | AF647 | 1/500 | Life Technologies | A-21449 | 1883471 |
| goat | mouse | CF680 | 1/500 | Biotium | 20065 | 14C0103 |
| goat | rabbit | CF660 | 1/500 | Bio-connect | 20369 | 14C0106 |
| goat | mouse | AF647 | 1/500 | Thermo Fisher Scientific | A-21236 | 2326487 |
| goat | rat | AF647 | 1/500 | Life Technologies | A-21247 | 1611119 |
| goat | mouse | CF660 | 1/500 | Bio-connect | 20368 | 14C0221 |

#### Imaging buffer and sample mounting

In this work we used two imaging buffers: a buffer with an oxygen scavenger (Glox-buffer) and a degassed buffer (N_2_-buffer). Glox-buffer was prepared as previously described [26]. Briefly, 1M stock solution of MEA (Sigma, 30070-10G, dissolved in 250 mM HCl) and glucose-oxidase plus catalase stock (70 mg/ml glucose-oxidase (Sigma, G2133-10KU, dissolved in Milli-Q), 4 mg/ml catalase (Sigma, C40-100MG) dissolved in Milli-Q) were prepared and stored at -80 °C. Just before imaging the final buffer was prepared by diluting MEA, glucose-oxidase plus catalase and glucose being in 50 mM Tris pH 8.0 (Final concentrations: 100mM MEA, 5% w/v glucose, 700 μg/ml glucose oxidase, 40 μg/ml catalase in 50mM Tris pH 8.0).

The N_2_-buffer uses a different method to remove oxygen from the imaging buffer [27]. A solution of 100 mM MEA in 50 mM Tris pH 8.0 was deoxygenated by smooth bubbling with N_2_ gas for 30 minutes using volumes 200-500 μl of buffer. The buffer was used immediately after this treatment.

Samples were mounted in closed off cavity slides (Sigma, BR475505) to prevent oxygen from entering the sample during imaging. The cavity slide was filled with approximately 90 μl of imaging buffer, after which the coverslip was flipped on top. Surplus buffer was removed from the sides of the coverslip using a vacuum pump to create a tight seal. Samples were used for up to an hour of imaging, because blinking behavior was compromised when imaging longer. Coverslip were removed and re-mounted in fresh buffer for a next round of imaging when necessary.

### Single-molecule detection and localization

Acquisitions were processed using the fast temporal median filter to remove constant fluorescence background [28]. Afterwards images were analyzed using the custom ImageJ plugin called DoM (Detection of Molecules, https://github.com/ekatrukha/DoM_Utrecht), which has been described in detail before [26]. Briefly, each image was convoluted with a combination of a Gaussian and Mexican hat kernel. By thresholding the images spots could be detected, after which their sub-pixel localization could be determined using an unweighted non-linear 2D gaussian fit of the original images using Levenberg-Marquardt optimization. Localizations with a width larger than 130% of the set detection PSF size were regarded as false positives. Reconstructions were generated by plotting each localization as a 2D Gaussian distribution with standard deviations in each dimension equal to the localization error. Drift correction was performed by calculating the spatial cross-correlation function between two intermediate reconstructions.

## Supporting information

Figure1_HighResolution

Figure2_HighResolution

Figure3_HighResolution

## Data availability

All data supporting the findings in this work are available at the Utrecht University online database Yoda. Source data are provided with this paper.

## Code availability

The custom code for analysis that support the findings of this work are available at the Utrecht University online database Yoda.

## Author Contributions

L.K and M.S. conceived research. M.S. developed the method presented in this work, with input from C.S.. Samples were prepared by D.J. and imaged by D.J. and M.S.. Data analysis was performed by M.S.. M.S., D.J. and L.K. wrote the manuscript with input from C.S.. L.K. supervised the project.

