## SupplementaryInformation for "3D multi-color far-red single-molecule localization microscopy with probability-based fluorophore classification"

### PSF-model and 2-channel MLE fit

The PSF in each channel is modelled as a 2D Gaussian where the intensity $\mu_{k,\lambda}$ at pixel location $k$ for the respective channel $\lambda$ is given by

$$\mu_{k,1} = \frac{\eta_1 N_{\mathrm{ph}} a^2}{2\pi\sigma^2} \exp\left[-\frac{\left(x_k - (x_0 - x_{\mathrm{align}})\right)^2 + \left(y_k - (y_0 - y_{\mathrm{align}})\right)^2}{2\sigma^2}\right] + b_1 \qquad 1$$

and

$$\mu_{k,2} = \frac{\eta_2 N_{\mathrm{ph}} a^2}{2\pi\sigma^2} \exp\left[-\frac{(x_k - x_0)^2 + (y_k - y_0)^2}{2\sigma^2}\right] + b_2 \qquad 2$$

with $N_{\mathrm{ph}}$ the total number of emitted photons, $a$ the pixelsize, $x_0$ and $y_0$ the position of the molecule, $x/y_{\mathrm{align}}$ a possible subpixel alignment correction between the channels, $\sigma$ the width of both Gaussian PSFs and $b_{1/2}$ the background in each channel. The two spots in channel 1 and 2 are fitted simultaneously, leading to 8 fit parameters ($\boldsymbol{\theta} = x_0, y_0, x_{\mathrm{align}}, y_{\mathrm{align}}, \sigma, N_{\mathrm{ph}}, b_1, b_2$).

The derivatives needed for the MLE fit routine with respect to the position are given by

$$\frac{\partial \mu_{k,\lambda}}{\partial x_0} = \frac{(x_k - x_0) N_{\mathrm{ph}} \eta_\lambda a^2}{2\pi\sigma^4} \exp\left[-\frac{(x_k - x_0)^2 + (y_k - y_0)^2}{2\sigma^2}\right] \qquad 3$$

and

$$\frac{\partial \mu_{k,\lambda}}{\partial y_0} = \frac{(y_k - y_0) N_{\mathrm{ph}} \eta_\lambda a^2}{2\pi\sigma^4} \exp\left[-\frac{(x_k - x_0)^2 + (y_k - y_0)^2}{2\sigma^2}\right] \qquad 4$$

where we dropped the channel alignment term. The derivative to $x_{\mathrm{align}}$ and $y_{\mathrm{align}}$ are similar to these derivatives for the first channel but zero for the second channel. The others derivatives are

$$\frac{\partial \mu_{k,\lambda}}{\partial \sigma} = N_{\mathrm{ph}} \eta_\lambda a^2 \frac{(x_k - x_0)^2 + (y_k - y_0)^2 - 2\sigma^2}{2\pi\sigma^5} \exp\left[-\frac{(x_k - x_0)^2 + (y_k - y_0)^2}{2\sigma^2}\right], \qquad 5$$

$$\frac{\partial \mu_{k,\lambda}}{\partial N_{\mathrm{ph}}} = \frac{\eta_\lambda a^2}{2\pi\sigma^2} \exp\left[-\frac{(x_k - x_0)^2 + (y_k - y_0)^2}{2\sigma^2}\right] \tag{6}$$

and

$$\frac{\partial \mu_{k,\lambda}}{\partial b_1} = \begin{matrix} 1 & \text{for } \lambda = 1 \\ 0 & \text{for } \lambda = 2 \end{matrix} \tag{7}$$

$$\frac{\partial \mu_{k,\lambda}}{\partial b_2} = \begin{matrix} 0 & \text{for } \lambda = 1 \\ 1 & \text{for } \lambda = 2 \end{matrix} \tag{8}$$

The log likelihood of the fit is given by

$$\log L = \sum_{\lambda=1,2} \sum_{k=x,y} \left(d_{k,\lambda} + \sigma_{\mathrm{noise}}^2\right) \log\left(\mu_{k,\lambda} + \sigma_{\mathrm{noise}}^2\right) - \mu_{k,\lambda} + \sigma_{\mathrm{noise}}^2 - \log\Gamma\left(\mu_{k,\lambda} + \sigma_{\mathrm{noise}}^2 + 1\right) \tag{9}$$

with $d_{k,\lambda}$ the observed pixel value and $\sigma_{\mathrm{noise}}$ the read noise of the camera [9]. In our setup we assume that the read noise of the EMCCD camera is zero. We use a Levenberg-Marquardt optimization routine to effectively maximize the log likelihood, where the next estimated fit parameters for iteration n+1 are given by

$$\boldsymbol{\theta}^{n+1} = \boldsymbol{\theta}^{n} + \left(\boldsymbol{H} + \gamma \operatorname{diag}(\boldsymbol{H})\right)^{-1} \boldsymbol{G} \tag{10}$$

with $\gamma$ the dampening factor tuned every iteration. The gradient $\boldsymbol{G}$ of the log likelihood is given by

$$G_i = \sum_{\lambda=1,2} \sum_{k=x,y} \frac{n_{k,\lambda} - \mu_{k,\lambda}}{\mu + \sigma_{\mathrm{noise}}} \frac{\partial \mu_{k,\lambda}}{\partial \theta_i} \tag{11}$$

and the Hessian $\boldsymbol{H}$ of the log likelihood is given by

$$H_{i,j} = -\sum_{\lambda=1,2} \sum_{k=x,y} \frac{n_{k,\lambda} + \mu_{k,\lambda}}{\mu + \sigma_{\mathrm{noise}}} \frac{\partial \mu_{k,\lambda}}{\partial \theta_i} \frac{\partial \mu_{k,\lambda}}{\partial \theta_j} \tag{12}$$

where the second derivatives with respect to the fit parameters are neglected as these are small near the optimum.

In the case for astigmatic 3D encoding we model the PSF as an asymmetric Gaussian given by

$$\mu_{k,1} = \frac{\eta_1 N_{\mathrm{ph}} a^2}{2\pi\sigma_x\sigma_y} \exp\left[-\frac{\left(x_k - (x_0 - x_{\mathrm{align}})\right)^2}{2\sigma_x^2} - \frac{\left(y_k - (y_0 - y_{\mathrm{align}})\right)^2}{2\sigma_y^2}\right] + b_1 \tag{13}$$

and

$$\mu_{k,2} = \frac{\eta_2 N_{\mathrm{ph}} a^2}{2\pi\sigma_x\sigma_y} \exp\left[-\frac{(x_k - x_0)^2}{2\sigma_x^2} - \frac{(y_k - y_0)^2}{2\sigma_y^2}\right] + b_1 \tag{14}$$

The derivatives with respect to the position are given by (again ignoring the alignment term)

$$\frac{\partial\mu_{k,\lambda}}{\partial x_0} = \frac{(x_k - x_0)N_{\mathrm{ph}}\eta_\lambda a^2}{2\pi\sigma_x^3\sigma_y} \exp\left[-\frac{(x_k - x_0)^2}{2\sigma_x^2} - \frac{(y_k - y_0)^2}{2\sigma_y^2}\right] \tag{15}$$

and

$$\frac{\partial\mu_{k,\lambda}}{\partial y_0} = \frac{(y_k - y_0)N_{\mathrm{ph}}\eta_\lambda a^2}{2\pi\sigma_x\sigma_y^3} \exp\left[-\frac{(x_k - x_0)^2}{2\sigma_x^2} - \frac{(y_k - y_0)^2}{2\sigma_y^2}\right] \tag{16}$$

The derivatives with respect to the widths of the Gaussian are given by

$$\frac{\partial\mu_{k,\lambda}}{\partial\sigma_x} = N_{\mathrm{ph}}\eta_\lambda a^2 \frac{(x_k - x_0)^2 - 2\sigma_x^2}{2\pi\sigma_x^4\sigma_y} \exp\left[-\frac{(x_k - x_0)^2 + (y_k - y_0)^2}{2\sigma^2}\right] \tag{17}$$

and

$$\frac{\partial\mu_{k,\lambda}}{\partial\sigma_y} = N_{\mathrm{ph}}\eta_\lambda a^2 \frac{(x_k - x_0)^2 - 2\sigma_y^2}{2\pi\sigma_x\sigma_y^4} \exp\left[-\frac{(x_k - x_0)^2 + (y_k - y_0)^2}{2\sigma^2}\right] \tag{18}$$

The other derivatives remain similar to the symmetric Gaussian.

Good initial estimates for the fit parameters are essential for finding the global optimum. The background was estimated as the median of the rim pixels of the ROI, the intensity as the total intensity of the background subtracted ROI and the position and width of the spot as the first and second moment of the background subtracted ROI respectively. To ensure physical results, the fit parameters were constrained; *x/y*-positions to [-200, 200] nm from center, $x/y_{\mathrm{align}}$ -positions to [-107, 107] nm, width to [100, 200] nm (2D) and [100, 400] nm (3D), intensity to [1 1e6] and background levels to [0 1e3].

### EM-CCD calibration

The EM-CCD was calibrated by imaging an out-of-focus knife-edge to create a smooth gradient over the FOV that covered the complete dynamic range of the camera. The gain and camera offset were then calibrated using the function *cal_readnoise()* from the DIPlib library in Matlab (https://diplib.org/). The gain of the camera scaled with the variance of the signal as $\mathrm{var}(S) = g\ \mathrm{mean}(S) + \sigma_{\mathrm{noise}}^2$ with $S$ the signal in ADU, $g$ the gain and $\sigma_{\mathrm{noise}}$ the read noise. The EM-CCD uses an electron multiplication process which drowns the read noise to zero ($\sigma_{\text{noise}} = 0$). However, this process introduces an additional noise called excess noise, which is indistinguishable from shot noise and effectively doubles it [29]. Therefore the true gain is overestimated by a factor of 2, but this allows the usage of a unmodified likelihood function (Equation 9).

**Supplementary Figure 1: Spectral characteristics of the setup**

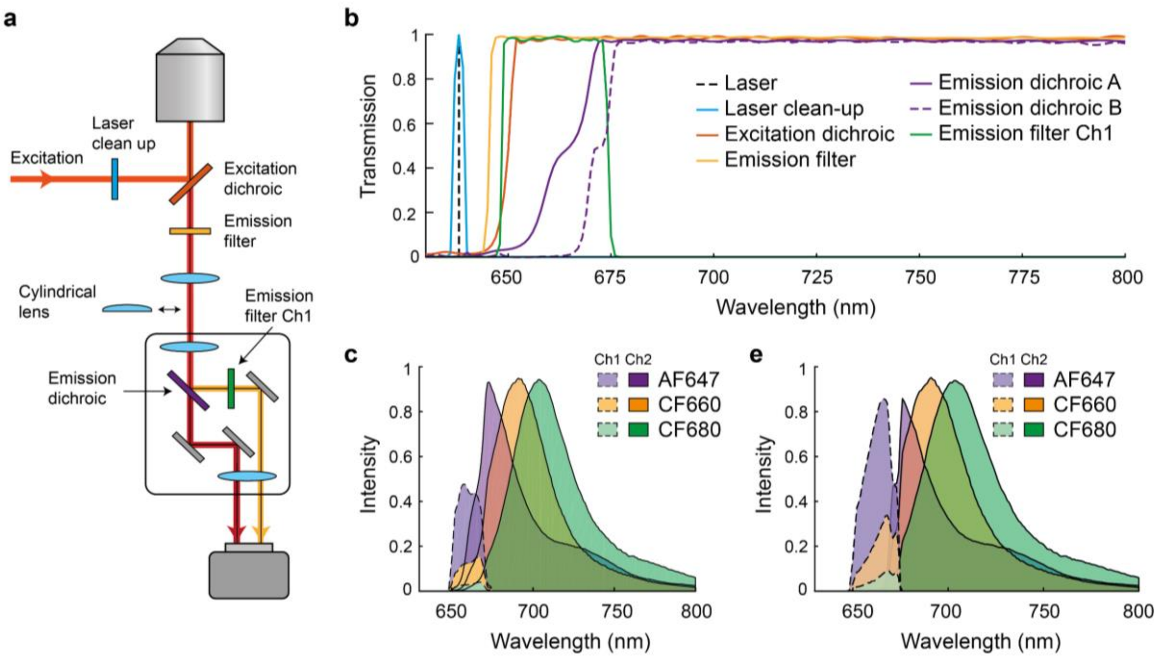


*a) Diagram of the setup. b) Spectral characteristics of all components. Emission dichroic A is used for 2-color imaging and emission dichroic B is used for 3-color imaging. c) Theoretical emission intensity in channel 1 and 2 of AF47, CF660 and CF680 with emission dichroic A. d) Theoretical emission intensity in channel 1 and 2 of AF47, CF660 and CF680 with emission dichroic B.*

**Supplementary Figure 2: Intensity histograms for 2-color and 3-color imaging**

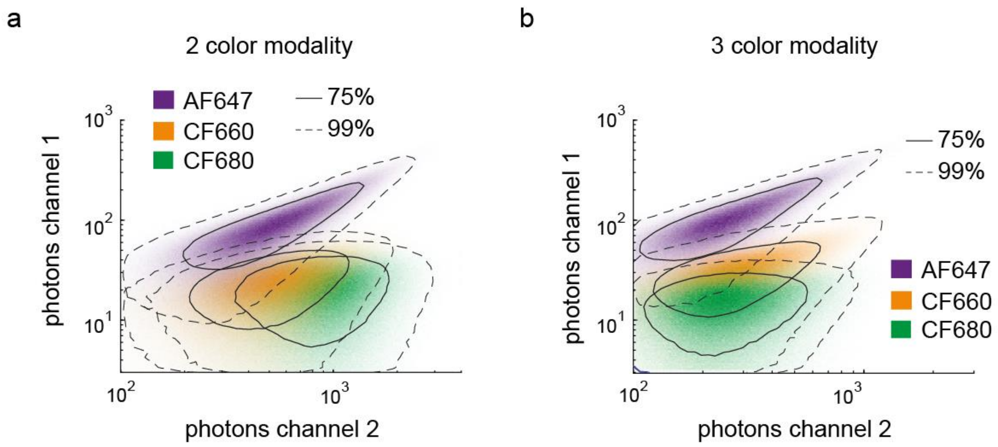


*2D histogram of intensity measured in channel 1 and 2 for 2-color (a) and 3-color (b) imaging for AF647, CF660 and CF680. Solid and dashed line indicate regions containing 75% and 99% of all events.*

**Supplementary Figure 3: Comparison between PFC and existing methods**

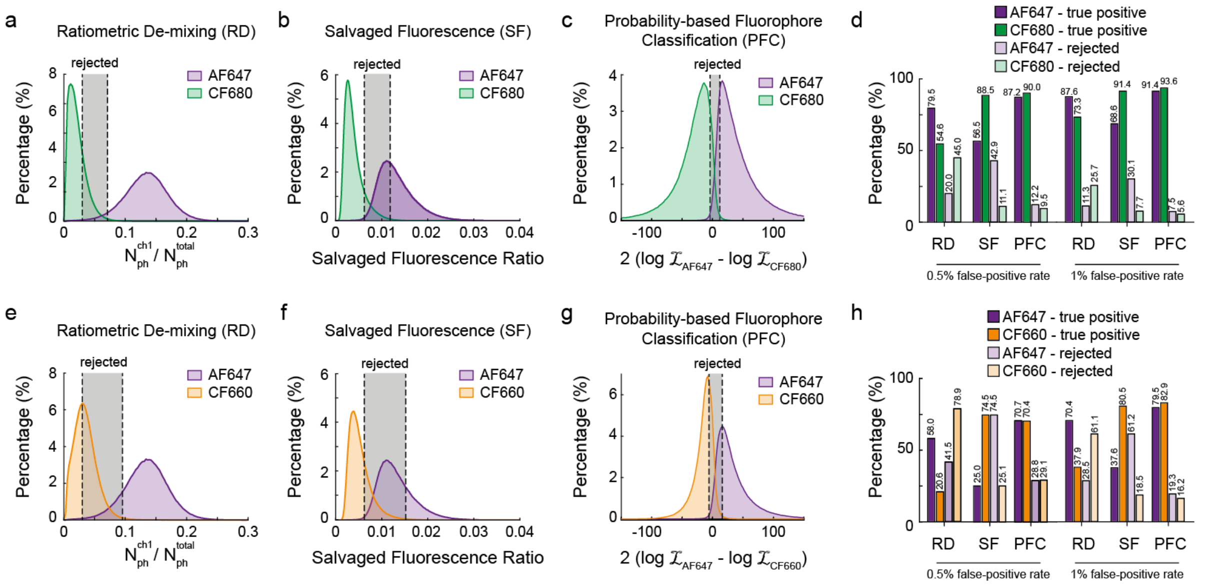


*a) Histogram intensity fraction in channel 1 for AF647 and CF680 when using Ratiometric De-mixing (RD). Grey area indicates the rejection zone to achieve 0.5% false-positive rate. b) Histogram of Salvaged Fluorescence Ratio (SF metric over intensity channel 2) of AF647 and CF680. Grey area indicates the rejection zone to achieve 0.5% false-positive rate. c) Histogram of the PFC GLRT of AF647 and CF680. Grey area indicates the rejection zone to achieve 0.5% false-positive rate. d) Classification performance for Ratiometric De-mixing (RD), Salvaged Fluorescence (SF) and PFC for false-positive rates of 0.5% and 1% with a photon threshold of 250. e-f) Same as (a-d) but for AF647 combined with CF660.*

**Supplementary Figure 4: Distribution of the GLRT for different photon counts.**

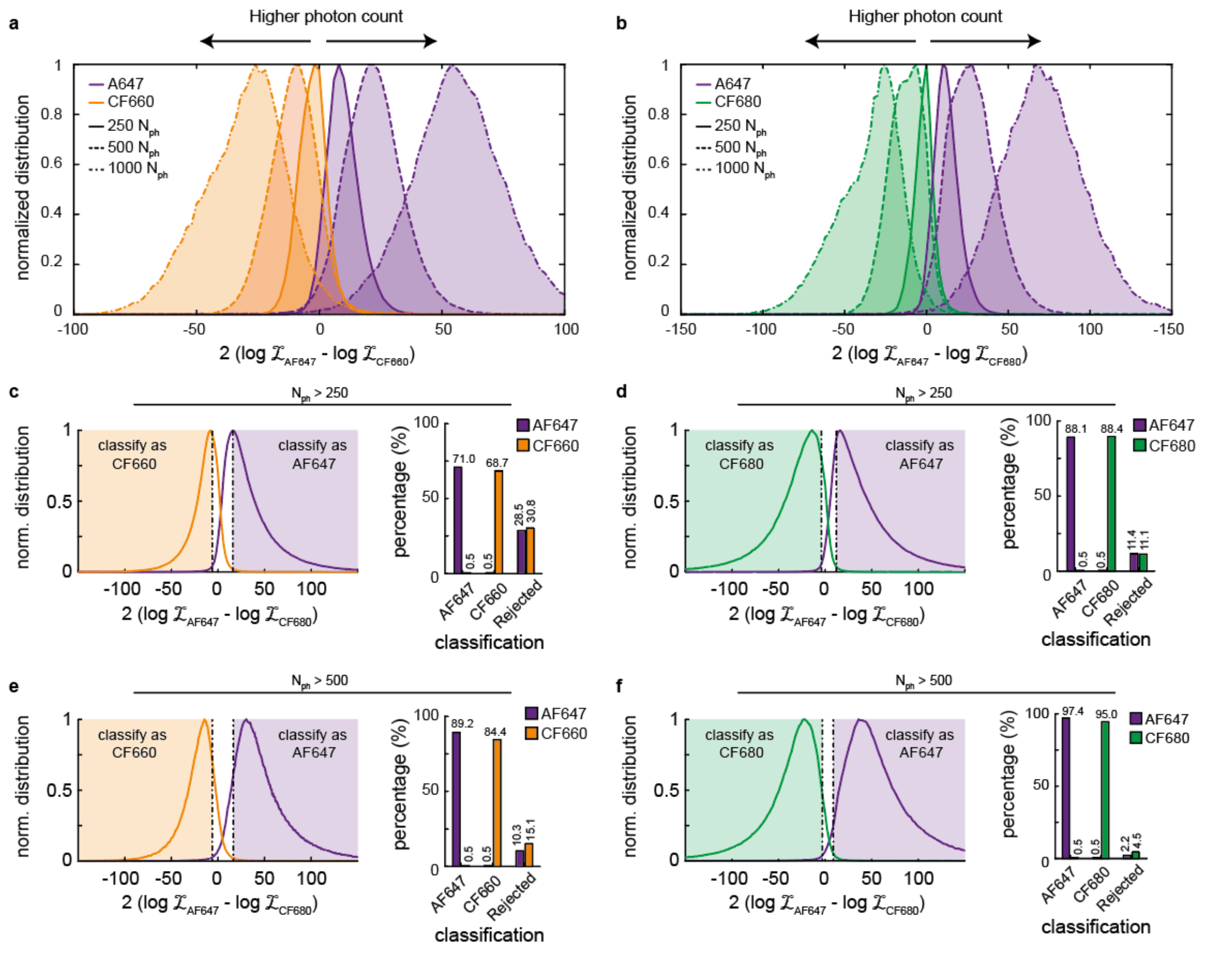


*a&b) Normalized GLRT distribution for AF647 vs CF660 (a) and AF647 vs CF680 (b) binned for different photon counts. c&d) Normalized GLRT distribution (left) and classification rates (right) for all events with a total intensity of 250 or more. e&f) Normalized GLRT distribution (left) and classification rates (right) for all events with a total intensity of 500 or more.*

**Supplementary Figure 5: Example PFC-dSTORM reconstructions.**

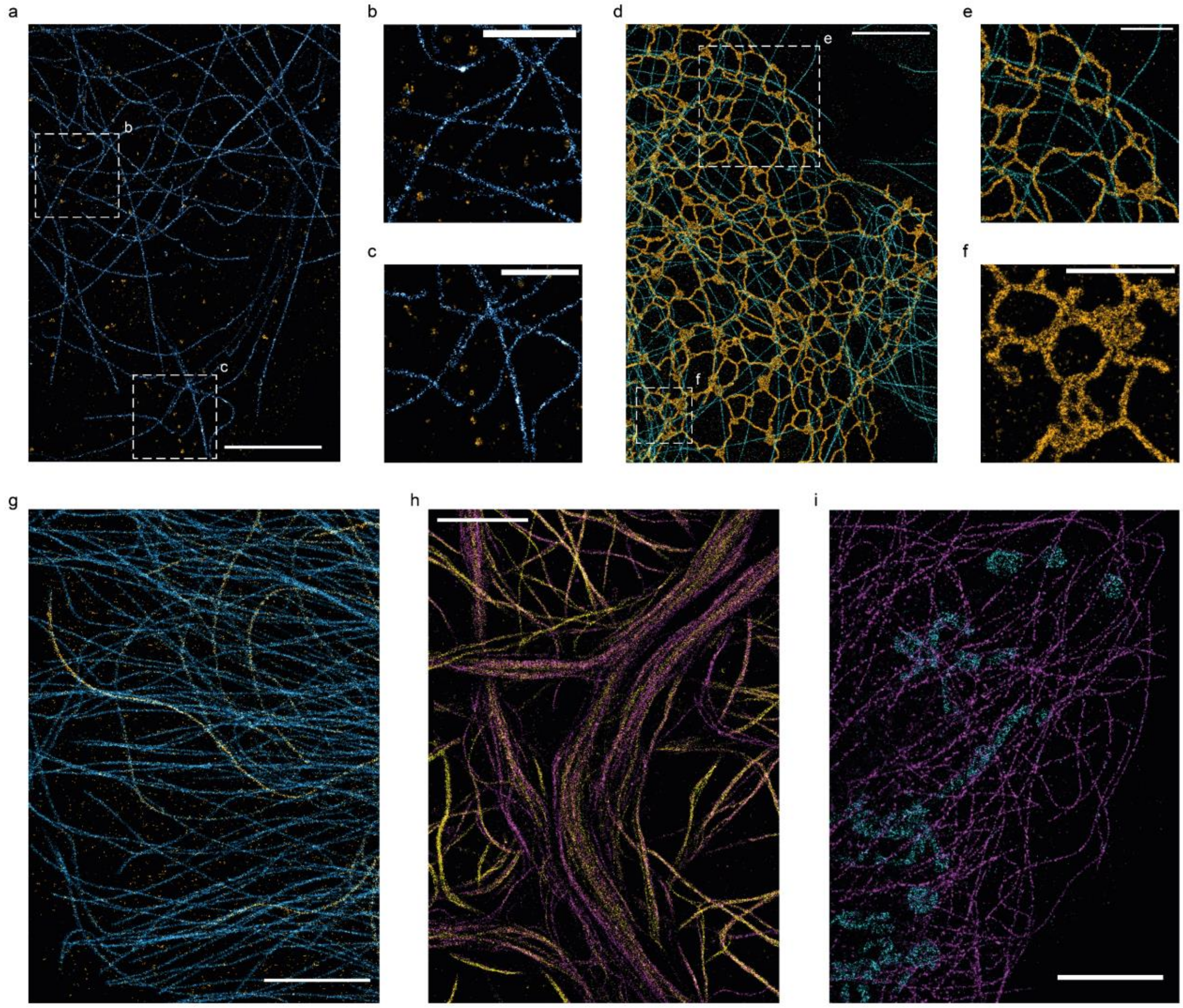


*a) COS-7 cell stained for alpha-tubulin (AF647, cyan) and clathrin HC (CF660, orange). b&c) Zooms of a. d) COS-7 cells stained for ER (SEC61b-GFP overexpression, AF647, orange) and alpha-tubulin (CF680, cyan). e&f) Zooms of c. g) U2OS cells stained for tyrosinated tubulin (AF647, cyan) and acetylated tubulin (CF660, yellow). h) Div14 neuron stained for tyrosinated tubulin (AF647, magenta) and acetylated tubulin (CF660, yellow). j) COS-7 cell stained for alpha-tubulin (AF647, magenta) and cytochrome C (CF680, cyan). Scale bars indicate 5 mm (a,d,g,h,j) and 2 mm (b,c,e,f).*

**Supplementary Figure 6: 2-color PFC-dSTORM on DIV14 neuronal synapses**

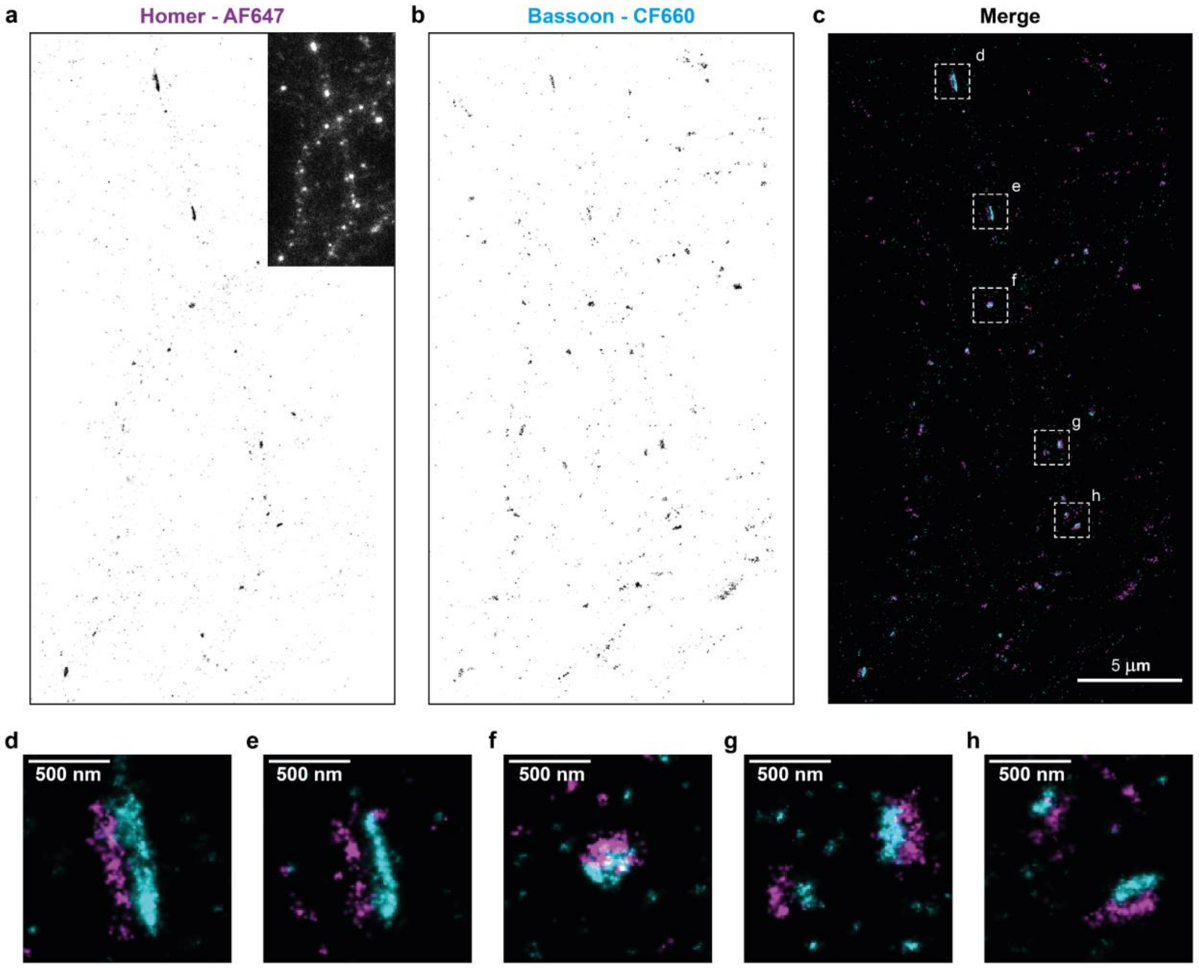


*a) Reconstruction of Homer (magenta) stained with AF647. Insert shows widefield image. b) Reconstruction of Bassoon (cyan) stained with CF660. c) Merge reconstruction of a&b. d-h) Zooms of c.*

**Supplementary Figure 7: Full FOV of image shown in Figure 2**

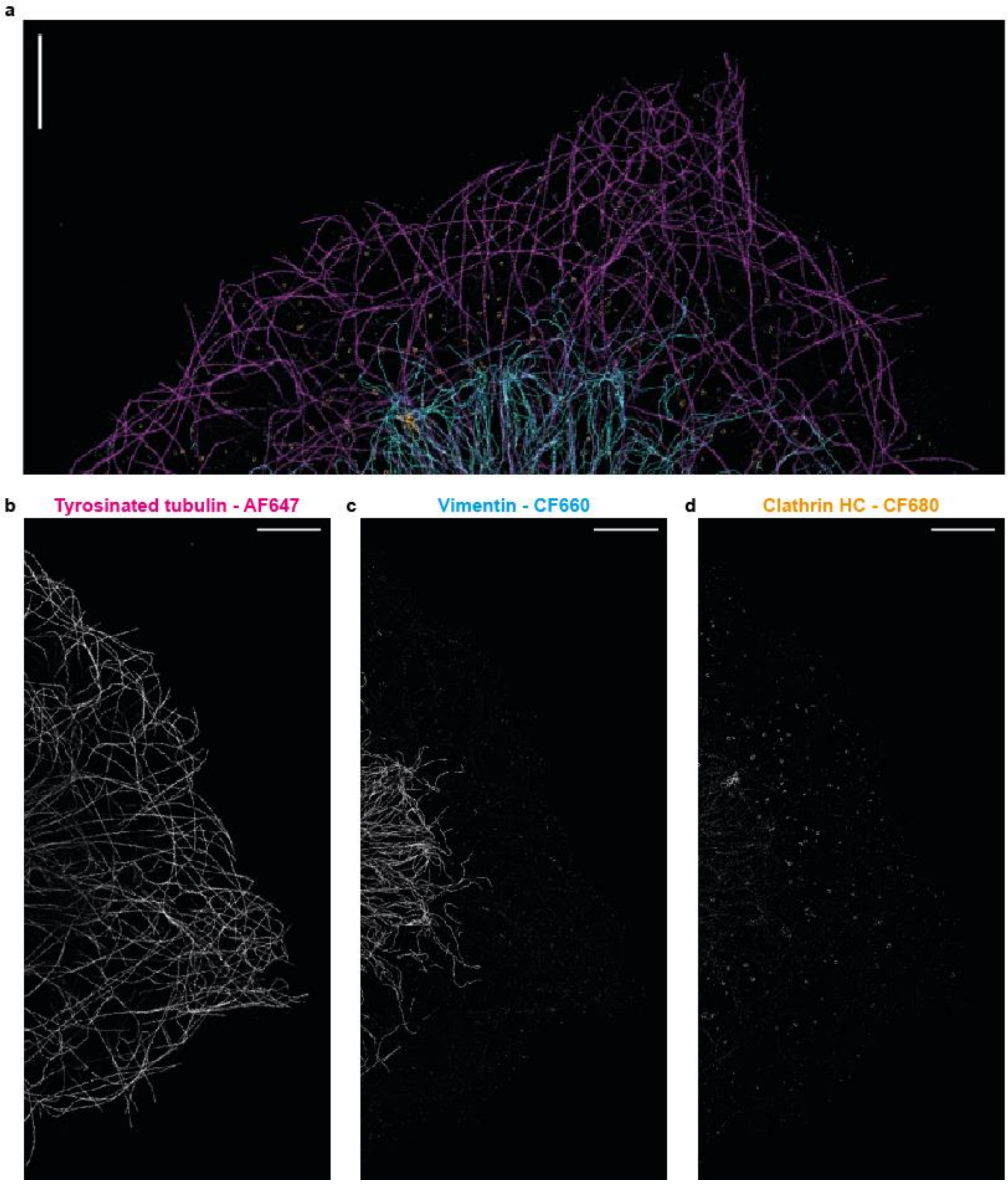


*a-d) 3-color PFC-dSTORM reconstruction of a COS7 cell stained for tyrosinated tubulin (AF647), vimentin (CF660) and Clathrin (CF680). a) Merge of all three channels. b-d) individual channels. All scale bars indicate 5 $\mu$m.*
