## Supplementary figures and images for "3D multi-color far-red single-molecule localization microscopy with probability-based fluorophore classification"

### Figure1_HighResolution

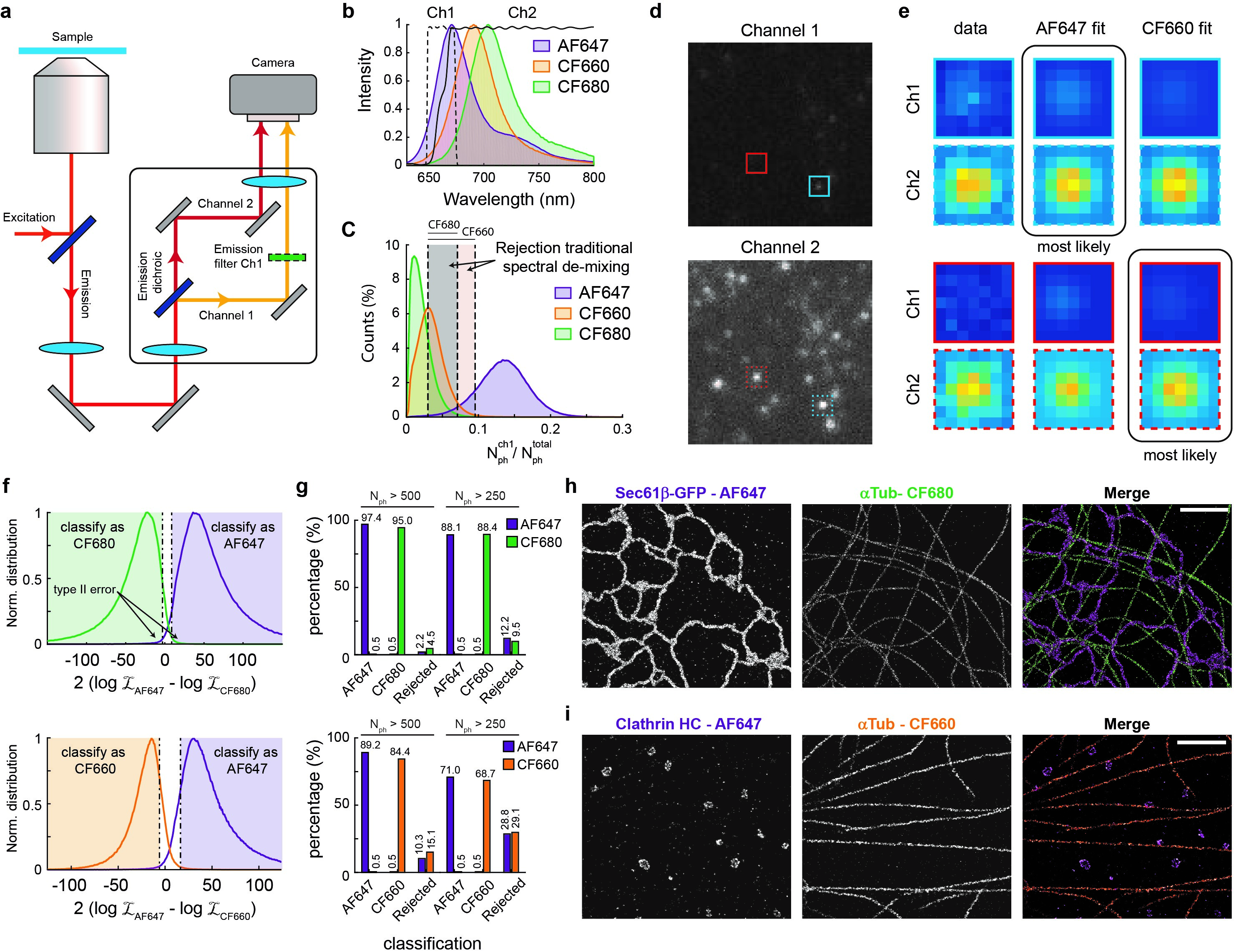

### Figure2_HighResolution

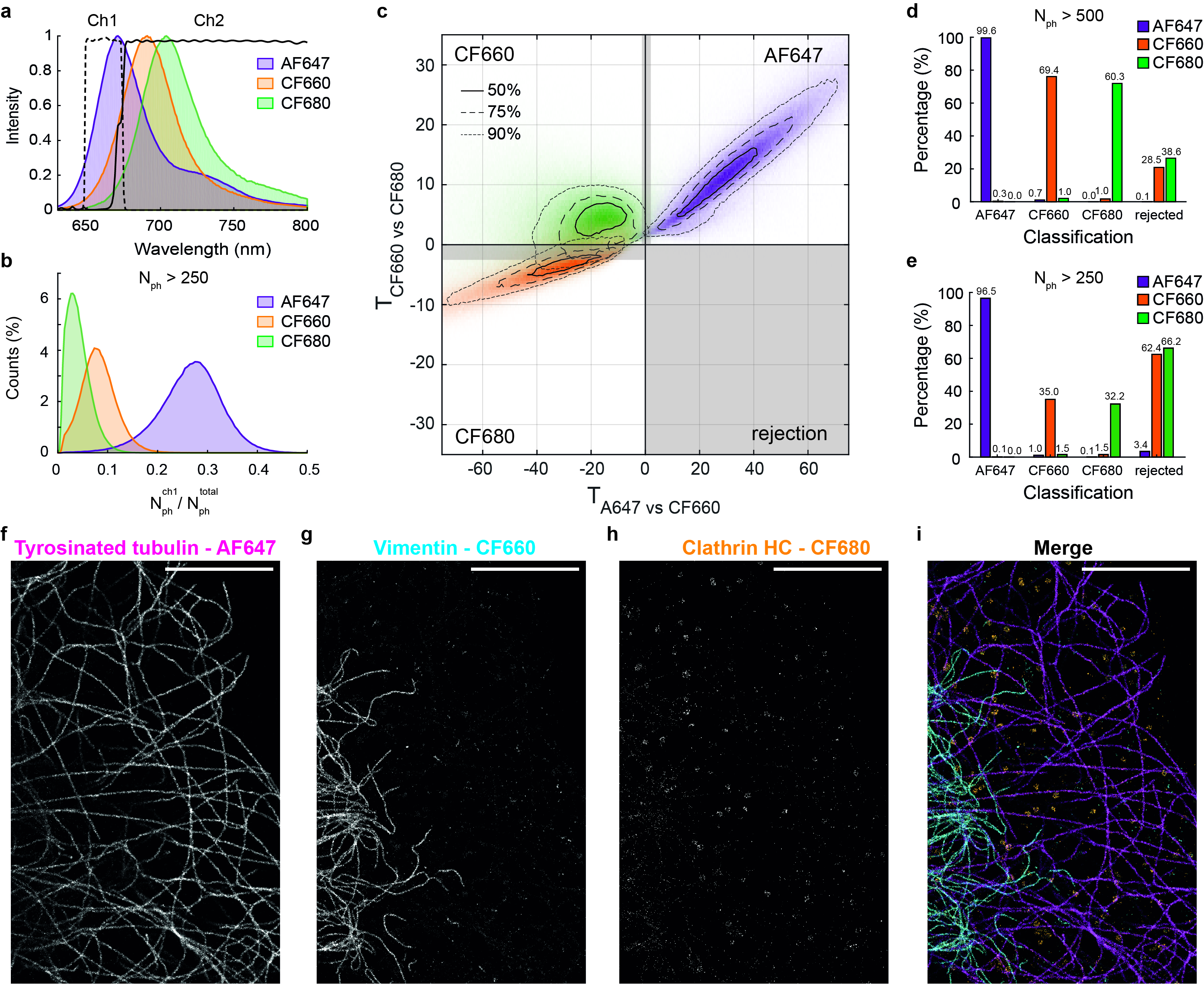

### Figure3_HighResolution

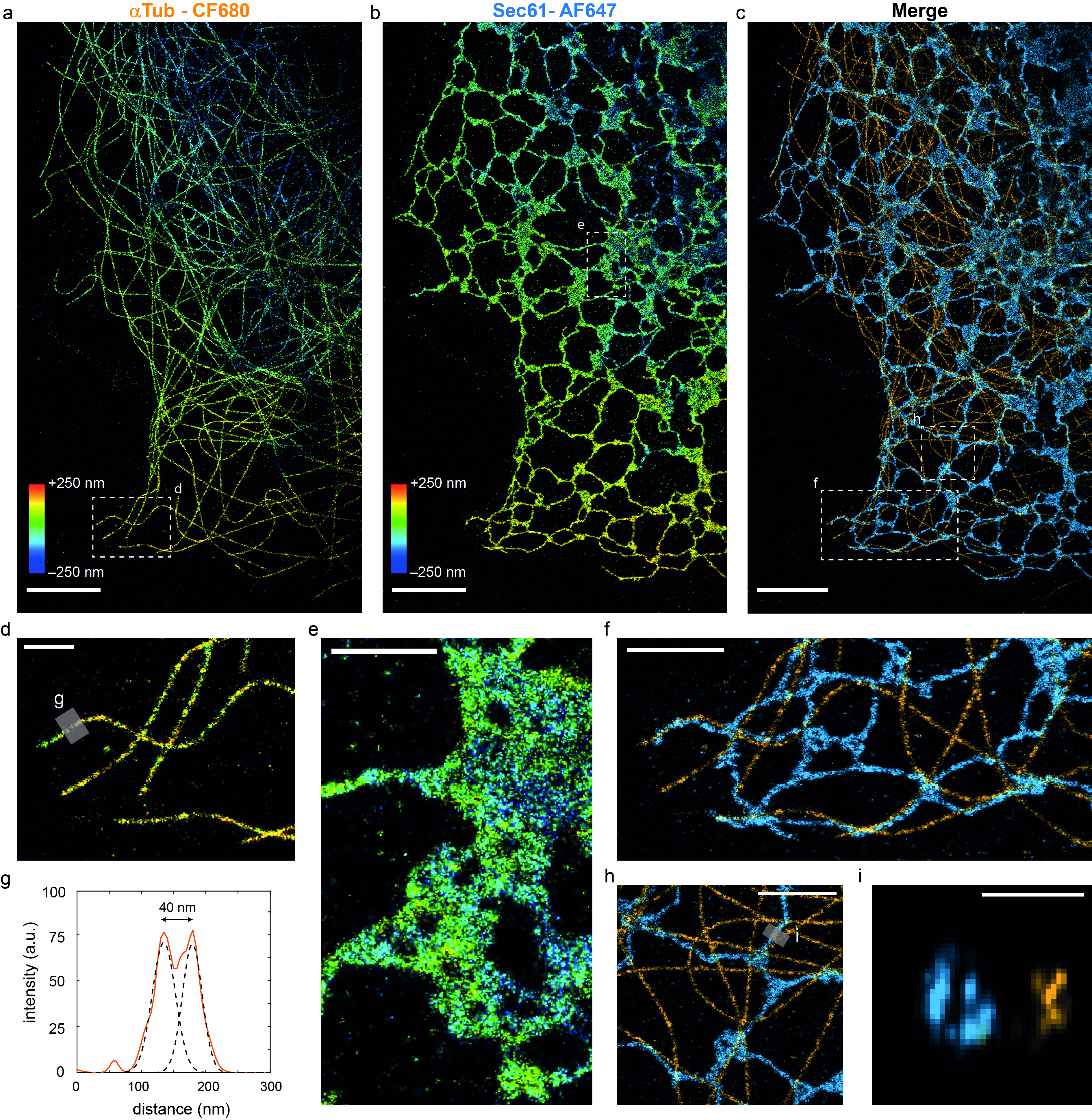
